# Behavioural and electrophysiological correlates of the cross-modal enhancement for unaware visual events

**DOI:** 10.1101/2023.12.01.569555

**Authors:** Márta Szabina Pápai, Mireia Torralba, Salvador Soto-Faraco

## Abstract

According to many reports, cross-modal interactions can lead to enhancement of visual perception, even when visual events appear below awareness. Yet, the mechanism underlying this cross-modal enhancement is still unclear. The present study addressed whether cross-modal integration based on bottom-up processing can break through the threshold of awareness. We used a binocular rivalry protocol, and measured ERP responses and perceptual switches time-locked to flashes, sounds or flash-sound co-occurrences. In behavior, perceptual switches happened the earliest when subthreshold flashes co-occurred with sounds. Yet, this cross-modal facilitation never surpassed the benchmark indicated by the probability summation, thus suggesting independence rather than integration of sensory signals. Likewise, the ERPs to audiovisual events did not differ from the summed unimodal ERPs, also suggesting that the cross-modal behavioural benefit for unaware visual events can be explained by the independent contribution of unisensory signals and suggest no need for a multisensory integration mechanism. Hence, even though cross-modal benefits appeared behaviourally, we suggest that this cross-modal facilitation might origin from well-known bottom-up attentional capture processes, contributed by each individual sensory stimulus.

## 1. Introduction

### 1.1. To what extent can MSI proceed in a purely bottom-up fashion, without any top-down regulation?

Our senses are constantly bombarded by inputs coming from different sensory modalities. Inferring which inputs belong together and which ones is not a non-trivial perceptual problem. It is often claimed (Driver & Spence, 2000; Stein & Meredith, 1993; Talsma & Woldorff, 2005) that this inference is guided by assumptions grounded on spatio-temporal correlations between events in different modalities, based on low-level (e.g. features) as well as high-level congruencies (meaning, learning associations). This cross-modal binding leads to multisensory integration (MSI) effects, which are claimed to carry beneficial consequences for information processing (Stein & Meredith, 1993). However, it is still controversial to what extent MSI can proceed in a purely bottom-up fashion, without any top-down regulation (Hartcher-O, Soto-Faraco, & Adam, 2017; ten Oever et al., 2016). This question is relevant because it bears on automaticity and, hence, on the limitation of processing resources required for MSI (Hartcher-O et al., 2017; ten Oever et al., 2016). Accordingly, a bottom-up MSI would show that multisensory interactions can take place under extremely impoverished sensory conditions. Indeed, such principle has been put forward as a means for rehabilitation in neurological patients suffering from visual hemineglect or hemianopia. Hence it is important to gain an understanding of the underlying mechanisms leading to such visual enhancement.

### 1.2. Multisensory integration below level of awareness with or without top-down regulation

Supporting a bottom-up view, a good number of recent studies have claimed that MSI can happen for events presented below awareness (Faivre, Mudrik, Schwartz, & Koch, 2014; Lunghi & Alais, 2013; Lunghi, Binda, & Morrone, 2010; Lunghi, Morrone, & Alais, 2014; Lunghi & Morrone, 2013; Zhou, Jiang, He, & Chen, 2010). Many of these studies have used binocular rivalry (BR), whereby a non-visual stimulus exerts an influence on a visual stimulus that is suppressed from awareness. However, the potential role of top-down mechanisms of attention or expectation related to time, space, or to other information provided by the non-visual stimulus (shape, orientation, semantics) are sometimes difficult to rule out (De Meo et al., 2015; Lippert, Logothetis, & Kayser, 2007). Nevertheless, a few studies have revealed MSI effects for stimuli below awareness, supposedly without top-down regulation (Lunghi & Alais, 2013; Lunghi et al., 2014; Zhou et al., 2010). Interestingly, in these studies cross-modal stimuli were typically presented for extended periods of time, promoting the build-up, or recall of existing associations between stimuli pairs that perhaps facilitated the cross-modal interactions below awareness (Mudrik, Faivre, & Koch, 2014). Therefore, the question remains as to whether MSI below awareness can happen when cross-modal stimuli do not share any feature-based, learned or semantic congruency, above and beyond mere spatio-temporal coincidence.

Recently, Pápai and Soto-Faraco (2017) addressed this question capitalizing on a BR protocol where cross-modal events resulted from the spurious co-occurrence of sounds and unaware flashes (on the suppressed eye) presented at unpredictable moments in time, and mutually uninformative. Despite perceptual BR switches happened earlier after cross-modal stimuli compared to unimodal (visual or auditory) events, suggesting an interaction between visual and sound signals (Miller, 1982), this benefit did not exceed statistical facilitation predicted by the probability summation model (PSM) (Otto & Mamassian, 2012; Raab, 1962). The most parsimonious explanation then, in line with the currently dominant models, was that the cross-modal benefit arose from the independent contribution of events in each modality, putatively involving bottom-up attentional capture (considering that even faint stimuli calls for bottom-up attention to a certain extent (Hancock & Andrews, 2007; Ooi & He, 1999; Paffen & Van der Stigchel, 2010). Thus, MSI between unaware visual events and sounds could not be concluded from these behavioural results. Yet, failure to violate the benchmark of the PSM does not negate MSI, by principle (Pannunzi et al., 2015), and therefore the initial question still remains unresolved.

### 1.3. The aim of the study: reveal the intertwined effect of MSI and attention

In fact, stimulus-driven effects in attention and MSI are often difficult to separate and deeply intertwined (Jon Driver & Spence, 1998; Macaluso, 2000; Mcdonald, Teder-Sälejärvi, & Ward, 2001). Considering that the distinction between attention and MSI behaviourally is quite challenging (i.e., since multisensory stimuli can integrate over quite wide spatial disparities (Bolognini, Rasi, & Ládavas, 2005; Frassinetti, Bolognini, & Làdavas, 2002; Spence & Driver, 1997; Teder-Sälejärvi, Russo, Mc Donald, & Hillyard, 2005) and within a flexible time window (Bolognini, Frassinetti, Serino, & Làdavas, 2005; Meredith, Nemitz, & Stein, 1987)), electrophysiological techniques might help provide evidence. Here, we gauge the possible neural correlates of bottom-up MSI effects using event-related potentials (ERPs) in a BR task akin to the one in Pápai & Soto-Faraco (2017). To the best of our knowledge, no previous studies have addressed the ERP correlates of audiovisual integration under binocular suppression of the visual event. Nevertheless, it is worth revising previous ERP findings regarding audiovisual integration for supra-threshold stimuli, as well as the correlates of unisensory visual events under suppression.

### 1.4. ERP findings of audiovisual integration

The neural correlates of MSI for supra-threshold stimuli are well established in the literature, and include expressions in sensory specific (Giard & Peronnet, 1999) as well as in multisensory brain regions (e.g., superior temporal sulcus, right insula, right prefrontal region) (Morís Fernández, Visser, Ventura-Campos, Ávila, & Soto-Faraco, 2015). Prior studies have reported that ERP responses to multisensory stimuli (modulation of visual evoked potentials by sounds or *vice versa*) often reveal nonlinearities, beyond the summed responses from the constituent unisensory stimuli (Fort, Delpuech, Pernier, & Giard, 2002a; Giard & Peronnet, 1999; Molholm et al., 2002; Talsma & Woldorff, 2005). These kinds of non-linearities (i.e., super-additive interactions) are interesting because they reveal MSI processes above and beyond additive effects. In particular, the auditory modulation on visual processing, of interest here, has been found to peak at occipital and parieto-occipital locations (Fort et al., 2002a; Giard & Peronnet, 1999; Molholm et al., 2002; Murray, Foxe, Higgins, Javitt, & Schroeder, 2001; Teder-Sälejärvi, McDonald, Di Russo, & Hillyard, 2002). What is more, some of these cross-modal effects occur very early after stimulus onset, indicating interactions within the first 200 ms (Fort, Delpuech, Pernier, & Giard, 2002b; Giard & Peronnet, 1999; Molholm et al., 2002), although the contribution of anticipatory processes cannot be ruled out (Talsma & Woldorff, 2005; Teder-Sälejärvi et al., 2002). Relevant for the purposes of the present study, Talsma and Woldorff (2005) tested for ERP correlates of cross-modal integration for simple audiovisual events presented at, or away from the focus of spatial attention. Remarkably, the results revealed that the ERP expression of MSI was attenuated for unattended events, compared to audiovisual events at attended locations. Along similar lines, another study from the same group showed that super-additive responses to cross-modal events diminish in absence of modality related top-down attention (Talsma, Doty, & Woldorff, 2007). In both cases, the attentional effect on MSI appeared at early time windows (within the first 200 ms), in line with the effect of attention on unisensory processing (Mangun, 1995; Woldorff et al., 1993). Therefore, considering the possible top-down effects at early time window (anticipation as well as attention) one might wonder to what extent were previously reported multisensory effects occurred independently of higher-order influences (Lippert et al., 2007; ten Oever et al., 2016). As far as MSI happens to occur above level of awareness it keeps being quite amenable to the modulation by higher-order mechanisms. However, because top-down influences can be more controlled (occasionally ruled out) below awareness (with a proper task), thus BR is a promising technique to single out bottom-up effects.

Previous findings about ERP correlates to visual events below awareness deserve some discussion here. Usually the P1, N1 and ‘late positivity’ (∼ P3) are the visual evoked potentials which are modulated as a function of awareness (Del Cul, Baillet, & Dehaene, 2007; Kaernbach, Schroger, Jacobsen, & Roeber, 1999; Koivisto et al., 2008; Koivisto, Revonsuo, & Lehtonen, 2006; Kornmeier & Bach, 2005; Marzi, Girelli, Miniussi, Smania, & Maravita, 2000; Mathewson, Gratton, Fabiani, Beck, & Ro, 2009; Metzger et al., 2017; Ojanen, Revonsuo, & Sams, 2003; Roeber et al., 2008; Valle-Inclán, Hackley, de Labra, & Alvarez, 1999). Generally, the finding about P1 and N1 is that their amplitude tends to diminish without awareness. Despite P1 effects have been suggested to simply reflect subjective visibility of the stimuli (Pins & Ffytche, 2003), there is more consensus about N1 visual potential, although the direction of the modulation by awareness remained controversial (reduced (Kaernbach et al., 1999; Koivisto et al., 2006; Marzi et al., 2000; Ojanen et al., 2003) or increased (Metzger et al., 2017; Valle-Inclán et al., 1999) N1 amplitude under suppression). Similarly, the late positivity component of the visual evoked potential is often modulated by awareness (Del Cul et al., 2007; Koivisto et al., 2008), but the interpretation of this modulation is less agreed upon (Eimer & Mazza, 2005; Koivisto et al., 2006).

### 1.5. The current study: to what and how?

In the current study, we used transient visual flash probes at threshold luminance embedded in rival gratings in a BR protocol. We decided for faint flashes to produce moderate stimulus-based perceptual switch, and leave some room for auditory modulation. Otherwise, the flashes occurred at unpredictable moments in time, and where intermixed with an unrelated, and also unpredictable sound. The two events would occasionally coincide. Both sensory events were task-irrelevant, thus mutually uninformative as well as unpredictable (in this way the anticipatory effects on the ERP were mitigated). Importantly, while top-down control is invariably at play in any task, including BR protocols, the present manipulation minimizes stimulus-selective top-down effects of attention or expectancy based on the temporal, spatial, feature-based or semantic congruency of the stimuli, within or across modalities. In this manner, we hoped to single out the effects of bottom-up signal processing, and their integration, if any. Since Pápai and Soto-Faraco (Pápai & Soto-Faraco, 2017) using very similar conditions found that behavioural correlates provided evidence for capture from each single modality but did not provide convincing proof of MSI thus, we expected to exploit the same logic here, and seek whether the neural responses (ERPs) to the stimuli allow concluding on MSI, given that they produce measurable unisensory responses. As mentioned earlier, the violation of the PSM in behaviour can be an indicator of the presence of MSI, although the lack of evidence for PSM violation does not rule MSI out (Murray et al., 2001; Pannunzi et al., 2015; Pápai & Soto-Faraco, 2017). Instead, MSI might be present, but express behaviourally below the threshold of PSM. Our approach here is to exploit such expression by using measurements of neural activity.

If MSI occurs for visual events below awareness, then we expect an amplitude modulation of the early visual evoked responses when combined with sounds, compared to the ERP responses to visual events alone, with special focus on parieto-occipital areas. We have chosen parieto-occipital areas to our region of interest, leading us to capitalize on the location of auditory modulation on visual processing (Giard & Peronnet, 1999; Molholm et al., 2002; Teder-Sälejärvi et al., 2002) and also on the location of the first stages of visual consciousness (Pins & Ffytche, 2003; Railo, Koivisto, & Revonsuo, 2011; Roeber et al., 2008). In order to produce the correct baseline, the ERPs to the visual alone and auditory alone stimulus conditions will be summed (hereafter referred to as the A+V ERP) and compared to the ERP response to the actual multisensory stimulus (AV ERP). Based on the principle of superposition of electrical fields, if we assume no MSI processes, then the AV ERP would be equivalent to the sum of the individual components, that is the A+V ERP. If the ERP to the multisensory event (AV ERP) deviates from the sum of individual ERPs (A+V ERP) then one should infer some integration process, in line with the co-activation model. However, in the contrast of multisensory versus sum of unisensory responses (AV-(A+V)) there would be double amount of baseline activity in the unisensory sum than in the multisensory response. In order to address this issue, activity without event presentation was taken into account, and subtracted from the unisensory sum (Talsma & Woldorff, 2005).

Furthermore, as we will measure behavioural responses (probability of perceptual switch in the BR task), likewise the pervious study of Pápai and Soto-Faraco (2017), we will be able to measure the probability of switch time-locked to the sensory events, and calculate whether suppressed audiovisual stimuli reveal the expected cross-modal facilitation in behaviour.

## 2. Methods

### 2.1. Participants

Data from 9 naïve observers was used (four female, average age 21.44 ± 2.1 years), and data from one additional participant was excluded (as he failed to run all of the blocks). The participants had normal or corrected-to-normal vision, normal stereo acuity, and presented no strong eye preference (as defined by perceptual predominance during binocular rivalry). The participants received 10 €/hour in return for taking part in the study.

### 2.2. Ethical statement

Participants gave written informed consent, and all methods were carried out in accordance with Declaration of Helsinki, under a protocol approved by the local ethics committee of the University of Pompeu Fabra (CEIC - Parc de Salut Mar).

### 2.3. Apparatus and Stimuli

Visual stimuli were created in MATLAB using PsychToolbox toolbox (Brainard, 1997; Pelli, 1997; Kleiner et al., 2007), displayed on a 19.8 inch CRT monitor (1024 x 768 pixels; 100 Hz refresh rate). The stimuli were displayed on a plain grey background (13.9 cd/m^2^). The two rival stimuli were contained within circular regions (11.5°) defined by a Gaussian envelope (SD= 0.13°). One rival stimulus was a horizontal Gabor grating with spatial frequency 1.2 cycles/° (mean luminance 17 cd/m^2^). The other rival stimulus was a radial checkerboard pattern whose mean luminance value was 19 cd/m^2^. Please note that the luminance values are the group average after the adaptive staircase measurement. This luminance imbalance was set to increase the suppression depth of the Gabor grating, which was always presented on the dominant eye. Both rival visual stimuli (Gabor patch, radial checkerboard) were black-and-white and low-contrast to favour multisensory integration (Jaekl, Pérez-Bellido, & Soto-Faraco, 2014; Perez-Bellido, Soto-Faraco, & Lopez-Moliner, 2013), and were centred on a black fixation cross (size of 0.25° and luminance of 3 cd/m^2^) surrounded by a grey circle in the centre (0.5°, luminance of 10 cd/m^2^). Additionally, each grating was surrounded by a black circle frame (0.2° width) presented simultaneously on the left and right halves of the monitor, with a centre-to-centre separation of 9.7°. The frames were binocularly matched therefore provided dichoptic stimuli for maintaining stable binocular alignment. These stimuli were viewed, one to each eye through a mirror stereoscope, giving a distance from the monitor to the eye of ∼57 cm. The observers’ head rested in a forehead-chin rest.

When subjects were exposed to the rival stimulus through the stereoscopically, they would experience alternations between the Gabor and the checkerboard. During this alternation, there were three types of event that could occur at random times; a visual flash presented on the lower part of the Gabor patch, a sound, or the flash and sound at the same time. The visual flash consisted of a 30 ms (10 ms fade-in/-out) contrast increment of the lower hemisphere of the Gabor grating. The size of the contrast increment was set individually to be at detection threshold under suppression (inter-participant average luminance =18 cd/m^2^). The sound was a 500 Hz, 40 dB, 30 ms tone (10 ms ramps in/out) with a 44.1 kHz sampling rate. The sounds where presented from two speakers located on the two sides of the monitor, vertically aligned to the location of lower half of Gabor grating. The timing of the audiovisual stimuli delivery occurred within 1 ms precision, as calibrated on a BlackBox Toolkit (Cambridge Research System). Responses were reported by key presses. The study was run in a dimly lit, sound attenuated test room.

### 2.4. Procedure

We required the participants to covertly monitor their current percept (Gabor patch or the radial checkerboard) meanwhile fixating on the fixation cross, by means of two keys. Participants were instructed to press both keys when a piecemeal mixture of the two patterns was visible. During the experiment, besides the planned pauses, participants were allowed to take a break any time they needed by releasing both of the keys. These trials were repeated in a subsequent run. Each observer participated in two or three 180-min experimental sessions, in two-three consecutive days (varied depending on the alternation rate of the subject).

#### 2.4.1. Pre-experiment calibration

After few minutes of dark adaptation, each experimental session began with several calibration runs, starting with the calibration of the mirror stereoscope, followed by a training period where participants became acquainted with the BR paradigm. Then, the relative dominance of the radial checkerboard was set between 65% and 75% using an up-down adaptive staircase to adjust contrast of the checkerboard (Kaernbach, 1990). Next, we set the individual threshold of probe detection for flashes on the Gabor patch under suppression. Please note, that probe threshold measured under suppression result in a relatively strong stimulus if presented under dominance (typical sensitivity lost in suppression is 0.3 to 0.5 log units relative to dominance (Lunghi et al., 2014; Nguyen, Freeman, & Wenderoth, 2001)). During threshold measurement, visual flashes were delivered during suppression periods with a delay of 1000 or 1500 ms locked to the initial of the suppression, and the contrast increments were adjusted by an adaptive staircase procedure designed to find the 50% detection threshold (Kaernbach, 1990). Finally, we tracked the natural alternation dynamics for a 3 minutes session in the very beginning of the experiment.

#### 2.4.2. Experimental run

The observers continuously monitored (and reported by key press) BR alternations between the Gabor patch and the radial checkerboard while their EEG were recorded. The visual, auditory or audiovisual events were presented at pseudo-random moments (see below), and each could be presented when the Gabor was reported dominant or suppressed; hence, the visual dominant (VD) or visual suppressed (VS) events, the auditory dominant (VD) or suppressed events, and the audiovisual dominant (AVD) or suppressed (AVS) events, respectively. Please note that dominant or suppressed, for the audio alone conditions, is a label variable denoting whether the Gabor grating was, respectively, dominant or suppressed, at the moment of sound presentation. This was done to align this condition with the other two in terms of perception. The intervals between events were composed of a fixed 5-s-delay plus a random 1-3 s jitter which was refreshed if a key press happened. Therefore, event presentation became temporally unpredictable and the events uncorrelated in time, which prevented possible top-down modulation and/or motor preparation (Teder-Sälejärvi et al., 2002) (**Fig 5**).

Since the events of interest were those under suppression, out of the 280 per modality condition, 3/4 were presented during Gabor suppression and 1/4 under Gabor dominance. Hence, each event modality was equally likely within each dominance condition. No events were ever presented during a piecemeal percept. Participants were asked to report binocular dynamics without any special instruction related to possible visual or audio abrupt stimuli. Considering the demanding nature of monitoring the stochastic fluctuation of rivalry and the unpredictive presentation time of the uninformative (i.e., task-irrelevant) events, we can assume little chance for task-related top-down systematic biases toward the stimuli, other than the introspective monitoring of the rivalry itself.

#### 2.4.3. EEG recording

During the experimental sessions, EEG data were acquired using 60 active electrodes (actiCAP, Brain Products GmbH, Munich, Germany) placed after the 10-20 international system, with the tip of the nose as online reference and AFz as ground. Data was referenced offline to the average of the left and right mastoids. Brain Vision Recorder (Brain Products, GmbH, Munich, Germany) was used for signal recording at a sampling rate of 500 Hz. Electrode impedance was kept below 5 kOhm. Horizontal (Heog) and vertical (Veog) electro-oculograms were recorded by two external electrodes, and used for off-line artefact rejection.

### 2.5. Analysis

#### 2.5.1. Behavioural data

In order to reduce inter-individual variability in overall alternation rate we normalized the absolute switch time by the natural alternation rate for each individual, measured in periods where no event was presented. We analyzed the time course of the probability of a switch in percept, time locked to event presentation, as a function of event type (time-probability analysis). This analysis was based on the probability of dominant/suppressed percept at each sampling point (0.025 alternation unit), and was informative as to how quickly the dominant percept changed after one of the events (presented under suppression/ dominance) in the design. One useful index in this type of analysis is the Mean Time to Switch (MTS), which indicated the time from event (A, V or AV under suppressed or dominant) presentation, when the probability of switch surpassed the probability of 50% of seeing that particular grating. Conscious report of the radial checkerboard was taken as the suppressed condition (the Gabor patch was not consciously perceived; the condition of interest), and conscious report of Gabor patch was taken as the dominant condition. Those trials where a switch happened before 250 ms were discarded from analysis, as the perceptual change was most likely not evoked by the event. We run a repeated measures ANOVA (with Greenhouse-Geisser correction where appropriate) on the MTS latency data with within participants’ factors: percept dominance (*Gabor dominant*, *Gabor suppressed*) and event modality (*audio*, *visual*, *audiovisual*), to compare switch latencies between modality conditions. The a priori hypothesis that audiovisual MTS would be shorter than unimodal ones was tested with one-tailed paired-t tests, whilst other contrasts were tested by two-tailed paired-t tests.

As part of our planned analysis, we also included a contrast of the probability of switch after audiovisual suppressed events against a PSM calculated from the unisensory suppressed events (Miller, 1982; Raab, 1962). The analysis has been run based on the adapted equation of probability summation for its use with BR switch times (Pápai & Soto-Faraco, 2017) (**Equation 1**).

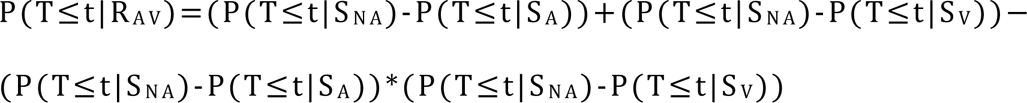

**Equation 1**. Let the probability of switch at time T after an audio event be P(T≤t|S_A_), and after a visual event be P(T≤t|S_V_), and the probability of the empirical distribution of natural alternation (of Gabor suppressed) P(T≤t|S_NA_). Then, one can model the theoretical audiovisual distribution based on the probability summation (redundant audiovisual: R_AV_), as the probability P (T ≤ t | R_AV_).

The equation considers the probability of switch after any of the unimodal events under suppression (P(T≤t|S_A_), P(T≤t|S_V_), after audio or visual events respectively), furthermore in order to avoid a double addition of variability unrelated to stimulus presentation, the probability distribution of natural alternation was also taken into account as Gabor suppressed P(T≤t|S_NA_). Then the modelled audiovisual distribution based on probability summation (P(T≤t|R_AV_) could be compared to the empirical probabilities distribution. The PSM provides a benchmark of what would be the quickest possible switch times that one would expect under the assumption of complete independence between visual and auditory sensory processing, according to a simple race model (Raab, 1962).

#### 2.5.2. ERP data analysis

The EEG data were filtered with a band pass filter between 0.5 and 25 Hz and an additional 50 Hz of notch filter was applied. Data from noisy electrodes were omitted, and the missing data was interpolated from the surrounding electrodes. EEG signal was segmented in epochs time-locked to the onset of suppressed visual, audio and audiovisual events ([-200 600] ms relative to the onset of the event). Additionally, we selected no-stimulus condition trials (used for ERP summation, see below) based on 800 ms pre-stimulus periods (from -1000 to -200 ms relative to the onset of suppressed or dominant events).

We performed automatic artefact rejection on resulting trials: all trials with amplitude exceeding ±100 μV in any of the scalp electrodes were excluded. Furthermore, the automatic artefact rejection was followed by a manual one, with special focus on the parietal and occipital electrodes. Thus, trials with eye-blinks, saccades, head movements, extensive muscle movements such as bite artefacts were removed, resulting in average 119 ±47 trials/condition after artefact rejection (from the 210 collected trials). Those trials where a switch happened before 250 ms were discarded from analysis, since those switches could not derived from the detection of visual flash.

Baseline activity was defined over the -200 ms to 0 ms period of each epoch for the three suppressed conditions. For the no-stimulus condition, we selected the first 200 ms of the trial. Event related potentials (ERPs) were computed for both the tree suppressed (audio, visual and audiovisual, respectively) conditions and the no stimulus condition, for each participant.

##### 2.5.2.1. Hypothesis-driven analysis

We focused on visual responses, in order to detect the possible effects of sound on visual processing, as a sign of early multisensory integration, partially in line with other studies (Fort et al., 2002a, 2002b; Giard & Peronnet, 1999; Teder-Sälejärvi et al., 2002). Therefore, we restricted our analysis to a visual region of interest (ROI). The ROI included the electrodes P3, P1, Pz, P2, P4, PO3, POz, PO4, O1, Oz, O2. ERP waveforms evoked by audio, visual or audiovisual events were analysed for this ROI area in the time window of 70-540 ms post-stimulus. In order to detect multisensory interactions, ERPs to the audiovisual events were compared to the algebraic sum of ERPs to the unisensory stimuli presented in isolation (i.e. audio and visual events), following earlier studies (Molholm et al., 2002; Molholm, Ritter, Javitt, & Foxe, 2004; Murray et al., 2001). Based on our hypothesis, differences between summed unisensory and the audiovisual ERPs, if any, would suggest nonlinear interaction. Yet, if the summed ERP responses from the unisensory presentations are equivalent to the audiovisual ERPs, then one would have to assume that independent neural responses to each of the unisensory stimulus are simply summed in the audiovisual event presentation (Murray et al., 2005). However, during the computation one needs to deal with the methodological problem of adding baseline activity twice, together with the actual ERPs, when calculating the sum of the individual responses (Talsma & Woldorff, 2005). Following Talsma & Woldorff (2005), to address this problem we calculated baseline neuronal activity what was present in absence of stimuli presentation or key press (no-stimulus condition). We used this no-stimulus activity as a mean to estimate the baseline EEG response and added it to the audiovisual responses before comparing it to the sum of audio and visual neural correlates. The audiovisual and summed audio and visual ERPs were compared using paired two-tailed t-test, and testing directional hypothesis one-tailed test (left), with significant differences being considered when they involved at least 18 consecutive data points (36 ms) at p<0.05. This criterion was decided following Guthrie and Buchwald approach (Guthrie & Buchwald, 1991), in order to correct multiple comparisons.

##### 2.5.2.2. Exploratory analysis

Beyond our main focus on the parietal-occipital areas, for completeness, we also looked at possible effects all over the scalp. We used paired t-tests (two-tailed, then one-tailed (right) to directional hypothesis) (significant level p=0.05) on all electrodes in the time window of 70-540 ms, to test the contrast between summed unisensory ERPs and audiovisual ERPs. ERP waveforms across conditions were compared with two-tailed paired t-tests. A cluster-based correction (Maris & Oostenveld, 2007) (10.000 randomizations) for electrodes and latencies was applied to correct for multiple comparisons. We used Fieldtrip toolbox (Oostenveld, Fries, Maris, & Schoffelen, 2011) and custom-made code for the exploratory statistical analysis.

## 3. Results

### 3.1. Behavioural results

#### 3.1.1. Time-Probability analysis

The probability of perceptual switches has been measured as a function of time, locked to the events. The moment of switches (Mean Time to Switch, MTS) has been indicated by the time when probability dropped down to 50%. The ANOVA on the latency data (MTS), returned a significant interaction between perceptual dominance (dominant, suppressed) and event modality (audio, visual and audiovisual) F(2,16)=3.891, p=0.042, which granted for further analysis. Supporting the presence of cross-modal effects, audiovisual events presented under suppression induced faster MTS (earlier emergence of the suppressed percept to awareness), compared to visual alone and audio alone events, one-tailed (left), paired t(8)=-2.775, p=0.012, and t(8)=-2.531, p=0.017 (α level after Bonferroni correction = 0.025). For completeness, we ran an additional two-tailed, paired, t-test (since we did not have a priori hypothesis concerning the direction of the difference) between visual and audio suppressed conditions, which were not different p=0.989. We ran a further two-tailed, paired t-tests between modalities under dominance, what failed to result in statistically significant effects (all p>0.05; **Fig. 1A-B**).

**Figure 1.**
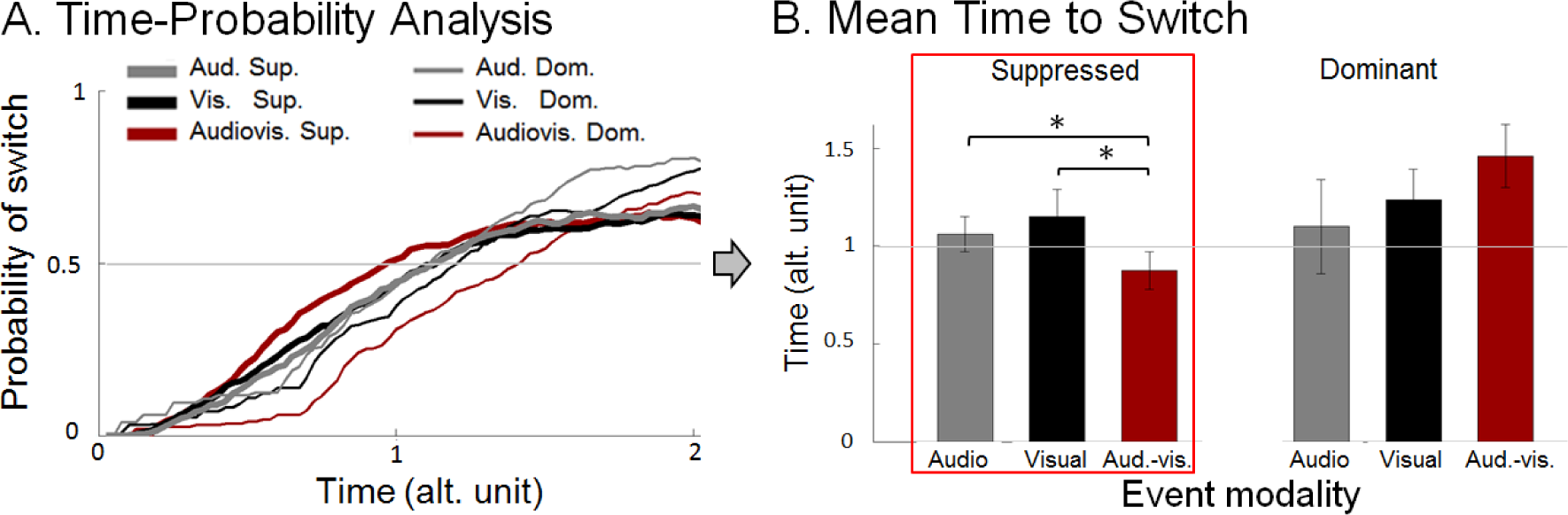
**A.** *Time-Probability Analysis.* The probability of switch is plotted as a function of relative time, expressed in alternation units (sampling points of 0.025 alt. unit). Switch in suppressed condition refers to change in perceptual state from seeing the radial checkerboard to seeing the Gabor patch (as events were presented on/during Gabor patch under suppression; audio, visual and audiovisual suppressed conditions, indicated by grey, black and red colours, respectively). On the other hand, switch in dominant conditions indicates change in perceptual state from seeing the Gabor to seeing the radial checkerboard (as events were presented on Gabor patch under dominance; audio, visual and audiovisual dominant). Zero time point indicates event onset time, which never occurred during piecemeal percept. **B.** *Mean Time to Switch (time at which probability crosses 50%*). The bars indicate the mean relative time of switch (in alternation unit with SEM) in each condition separately. Additionally, please note that significant comparisons marked by ‘*’.

#### 3.1.2. Probability summation

Additionally, we tested the empirical behavioural data from the audiovisual suppressed condition against the PSM calculated from the unimodal suppressed conditions and the natural alternation rate (see Methods). The statistical comparison between the PSM and the empirical AV suppressed data throughout time did not reveal violations of the model (one-tailed (right), paired, t-test, all p>0.05). Thus, the data clearly illustrates a facilitation of audiovisual events under suppression compared to unisensory conditions, but not a violation beyond the race model (this replicates the behavioural study of Pápai and Soto-Faraco (Pápai & Soto-Faraco, 2017)).

### 3.2. ERP results

#### 3.2.1. Hypothesis-driven analysis: Multisensory ERP responses at occipito-parietal electrodes

Mean ERP values of the ROI electrodes were calculated for audio, visual and audiovisual events. We tested for differences between the summed auditory and visual ERPs and the audiovisual ERPs (summed to the no-stimulus ERPs, see Methods). The paired t-tests (two-tailed) in the time window of 70-540 ms did not reveal any significant deviance (after applying Guthrie and Buchwald’s correction for multiple comparisons (Guthrie & Buchwald, 1991) (**Fig. 2**)). Please note this contrast is mostly frequently done using a two-tailed test, since it is more conservative and hence, any significant effects detected can be interpreted more confidently (Molholm et al., 2002; Murray et al., 2001; Teder-Sälejärvi et al., 2002). We followed the same tradition here. However, given this first negative finding, in order to increase sensitivity to multisensory effects if any, we lowered the significance threshold assuming a one-tailed (directional) paired t-test, based on the assumption of multisensory ERP reveals in higher amplitude than the unisensory sum. This comparison suggested a small window of significant difference after correction, but late in the ERP (between 394-432ms). This effect then happened after early sensory processes. Hence, like the behavioural responses, the ERPs did not reveal a convincing sign of genuine bottom-up MSI between sounds and unaware visual events. Still, in order to understand the underlying mechanisms further, we decided to run some complementary analyses.

**Figure 2.**
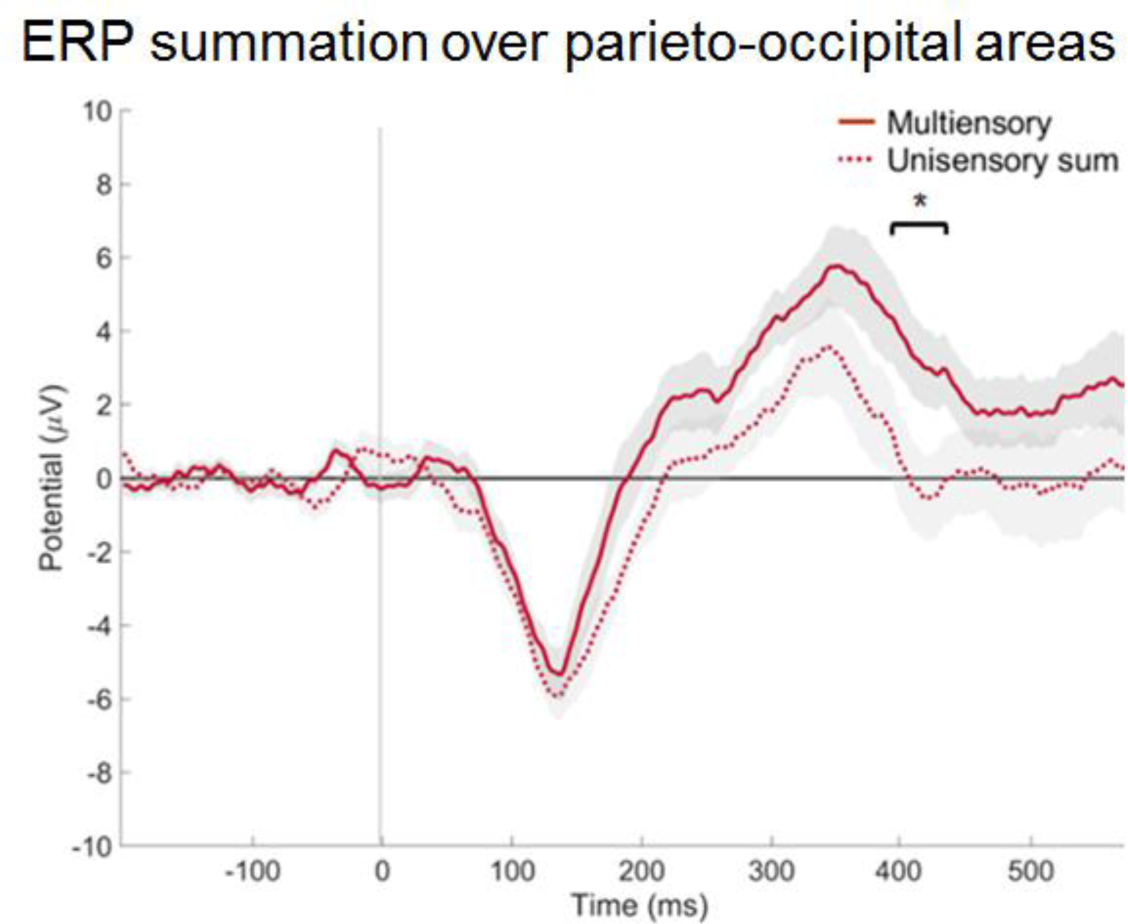
*ERP summation over parieto-occipital areas.* The summed audio and visual suppressed ERPs, called ‘Unisensory sum’ is contrasted against the sum of audiovisual suppressed and no-stimulus ERPs, called ‘Multisensory’ over the average of ROI areas (P3, P1, Pz, P2, P4, PO3, POz, PO4, O1, Oz, O2) with SEM. The paired t-test (one-tailed (right)) suggested some statistically significant difference, in the time window of 394-432ms after Guthrie and Buchwald’s correction. The 0 time point corresponds to stimulus onset.

#### 3.2.2. Visual evoked potentials (VEPs) over ROI area

Here, we addressed the effects of awareness on VEPs, to check if the visual events under suppression generated a measurable response and also, if our results were in line with previous studies measuring ERPs to visual events presented below awareness. Because of the 3:1 difference in suppressed and dominant number of trials in our design, we randomly picked trials from the suppressed condition to equate the less populated dominant condition (40 +-7 trials/condition after artefact rejection). We then calculated the average VEPs over the ROI area, for suppressed and dominant conditions (**Fig. 3A**), and ran paired t-test (two-tailed) between the two waveforms within the time window 70-540 ms. The P1 waves were not very pronounced, what is maybe not so surprising considering the weak stimulus strength of the visual flash (Pins & Ffytche, 2003). Additionally, both the suppressed and dominant VEPs showed N1 and late positivity (∼P3) visual responses. The N1 response to dominant visual events appeared larger than the suppressed one, in line with many previous studies (Kaernbach et al., 1999; Koivisto et al., 2006; Marzi et al., 2000; Ojanen et al., 2003), and late positivity was reduced in suppressed compared to dominant condition, again, in line with what has been shown in the past (Kaernbach et al., 1999; Railo et al., 2011; Roeber et al., 2008) (despite some discrepancies exist for both of the components (Metzger et al., 2017; Valle-Inclán et al., 1999)). Yet, please note that these differences, albeit mostly being in the expected direction, did not reflect statistical significance after Guthrie and Buchwald’s correction (all p>0.05). Still, the relevant result here is that the VEP appeared for suppressed visual events, whether weaker than the dominant ones or not, but ensuring that these ERPs could be indeed measured in our paradigm.

**Figure 3.**
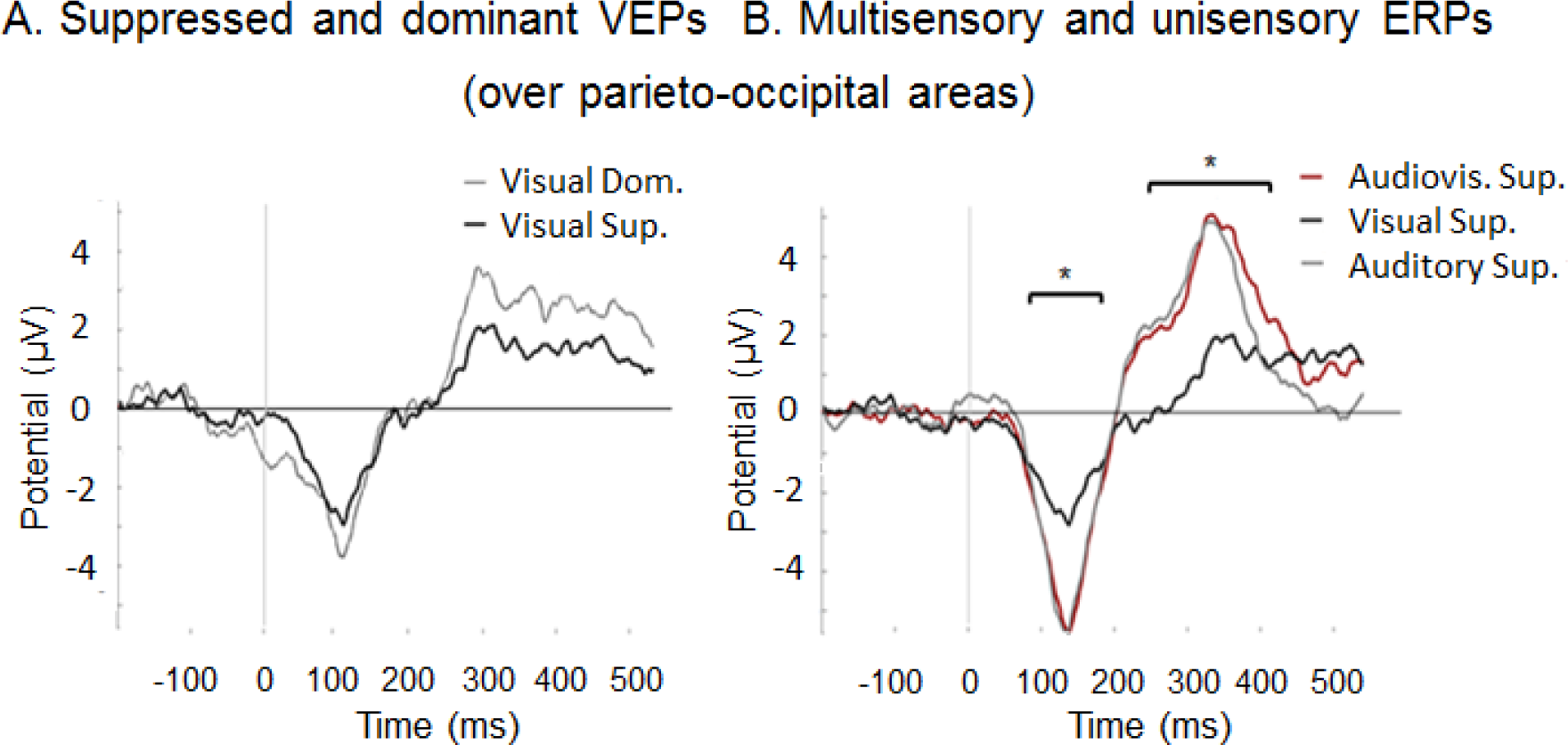
**A.** *Suppressed and dominant VEPs.* Visual N1 and late positivity (∼P3) over ROI (P3, P1, Pz, P2, P4, PO3, POz, PO4, O1, Oz, O2) areas evoked by dominant or suppressed visual event are plotted respectively. There was no statistically significant difference found between the conditions. Visual dominant condition is indicated by grey, the visual suppressed by black colour. **B.** *ERPs of multisensory and unisensory conditions.* ERP are presented in audiovisual, visual and auditory suppressed conditions, marked by red, black and grey colours, respectively over ROI areas. Significant differences between visual and audiovisual suppressed conditions are marked by ‘*’. In both of the graphs 0 time point corresponds to stimulus onset time.

#### 3.2.3. Multi- and unisensory evoked potentials over ROI area

The comparison of audiovisual ERPs to the summation of unimodal ERPs presented above did not reveal convincing signs of clearly bottom-up MSI, very much like the behavioral effects did not violate the PSM benchmark. Still, in order to further understand the results, we inspected the single modality and audiovisual ERPs over the ROI area. The paired t-tests (two-tailed) ran in the time window 70-540 ms after the event revealed statistically significant differences between audiovisual and visual ERPs (92-166 ms and 208-432 ms, all p<0.05, corrected). Remarkably, the audiovisual and audio ERP waves were not different (all p>0.05), in fact they seemed to overlap quite a lot (**Fig. 3B**). This pattern of results indicated that the ERP response to the audiovisual events was clearly dominated by the auditory evoked potential. This was true despite there was a visual response when measured alone, and that the electrode cluster we focused on should be visually responsive. Thus, the behavioral advantage in switch latencies for audiovisual events was not particularly reflected in the ERP responses over this ROI and time window. This is perhaps not surprising, because of the nature of the behavioral effects does not lead to conclude on a co-activation, or non-linear interaction at the sensory level.

#### 3.2.4. Exploratory analysis: Multisensory ERP responses over all of the scalp

According to the hypothesis-driven ROI analyses above, MSI cannot be concluded based on the ERP response. One assumption underlying those analyses, based on a good number of previous papers, was that MSI effects might express as a modulation of the VEP, over the parieto-occipital areas, hence the use of an ROI. Still, one might raise the question that putative multisensory effects might express at other scalp locations. Here, we ran exploratory analyses on the ERPs across the scalp. We used the electrode-by-electrode average data over all the scalp locations for each condition (**Fig. 4A**) in the time window of 70-540 ms. We first ran two-tailed, followed by a one-tailed (right) paired t-test in order to reveal any weak multisensory effect, if any, by using for the same contrast used in the ROI analysis: comparing the summed unisensory ERPs (audio suppressed plus visual suppressed) versus audiovisual ERPs plus the no-stimulus ERPs (**Fig. 4B**). After, cluster-based correction, there were no significant differences (all p>0.05). For completeness, similar to the ROI analysis, we also tested for differences between each unisensory and the audiovisual ERPs, at each electrode/time (by paired, two-tailed t-tests). Similarly to the ROI analysis, the statistical test revealed significant differences between audiovisual and visual conditions ERPs: 99-184 ms negative ERP shift over centro-medial and parietal areas all p<0.05 (corrected), a positive shift 192-426 ms over cento-medial areas, all p<0.05 (corrected), and a later negative shift 442-540 ms over centro-medial areas, all p<0.05 (corrected) (**Fig. 4**). Yet, the audiovisual ERPs did not differ from auditory ERPs. Thus, all in all, the pattern resulting from the analysis across the scalp was very similar to the one over the ROI area: the summed unimodal ERPs accounted for the audiovisual ERPs.

**Figure 4.**
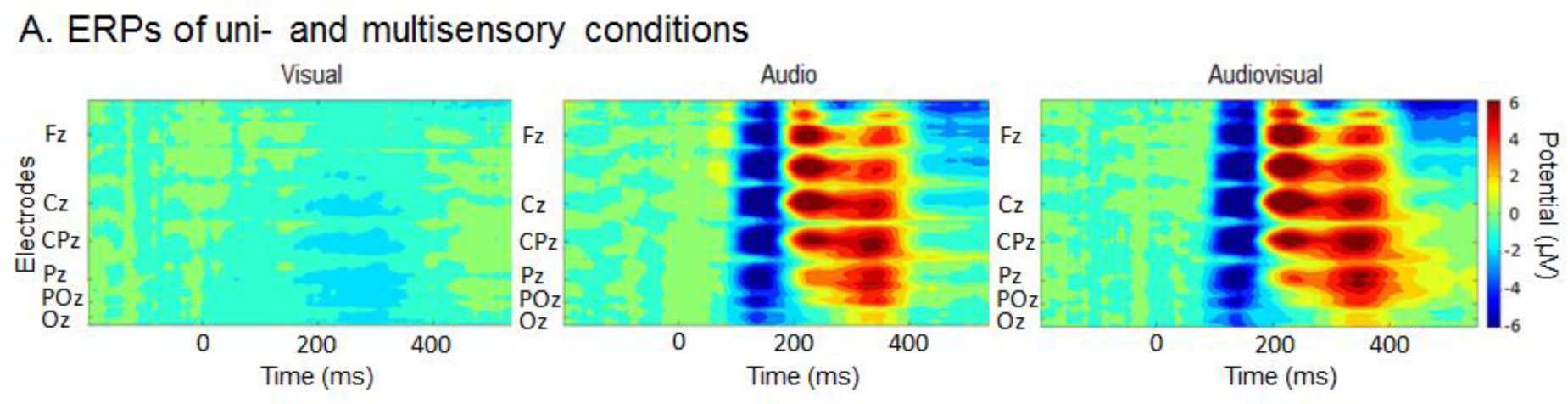
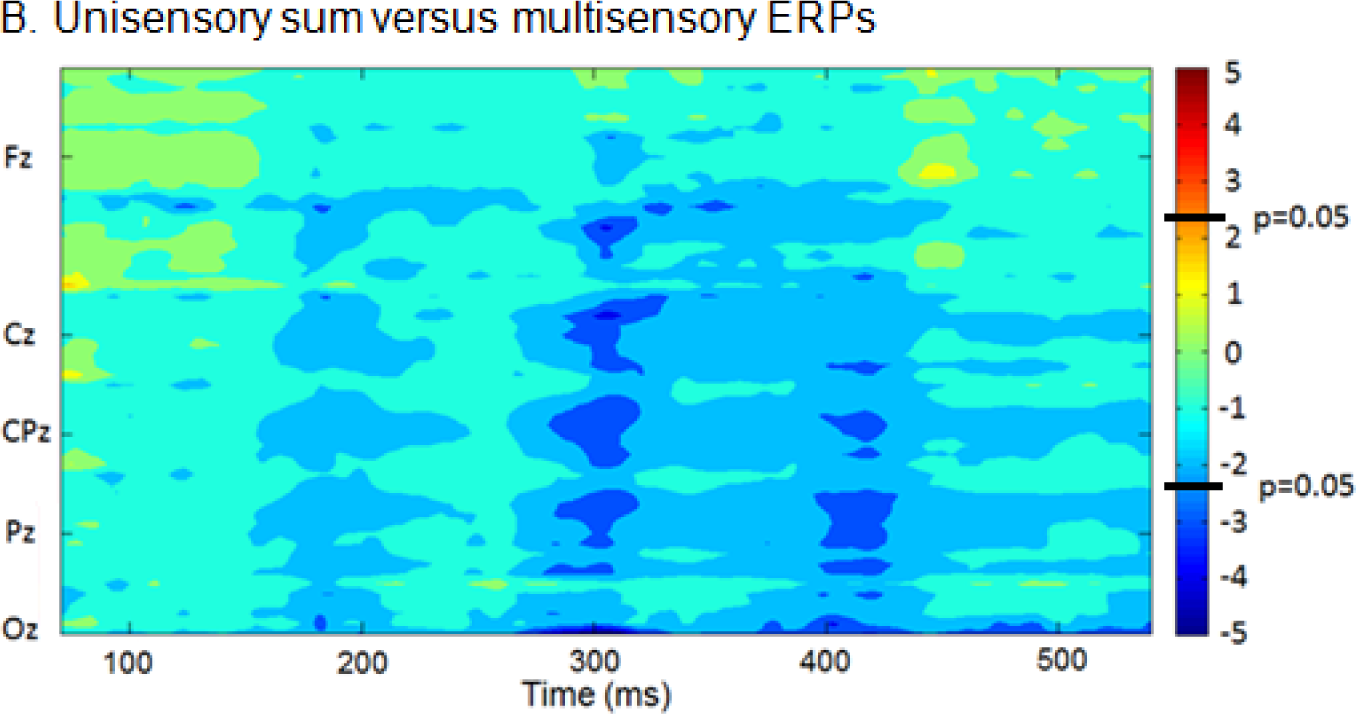
**A.** *ERPs of uni- and multisensory conditions.* ERP potentials (µV) are presented in visual, audio and audiovisual conditions as a function of time in all electrodes. 0 time point corresponds to stimulus onset time. **B.** *Unisensory sum versus multisensory ERPs.* The t-values of the paired t-test (two-tailed) are plotted as a function of time (70-540ms), across the scalp. The paired t-test did not reveal any statistically significant difference of this contrast after Guthrie and Buchwald’s correction. Colour map shows t-values, from positive values (red) to negative values (blue). Please note that the t-values of ±2.2622 correspond to the p-value of 0.05.

**Figure 5.**
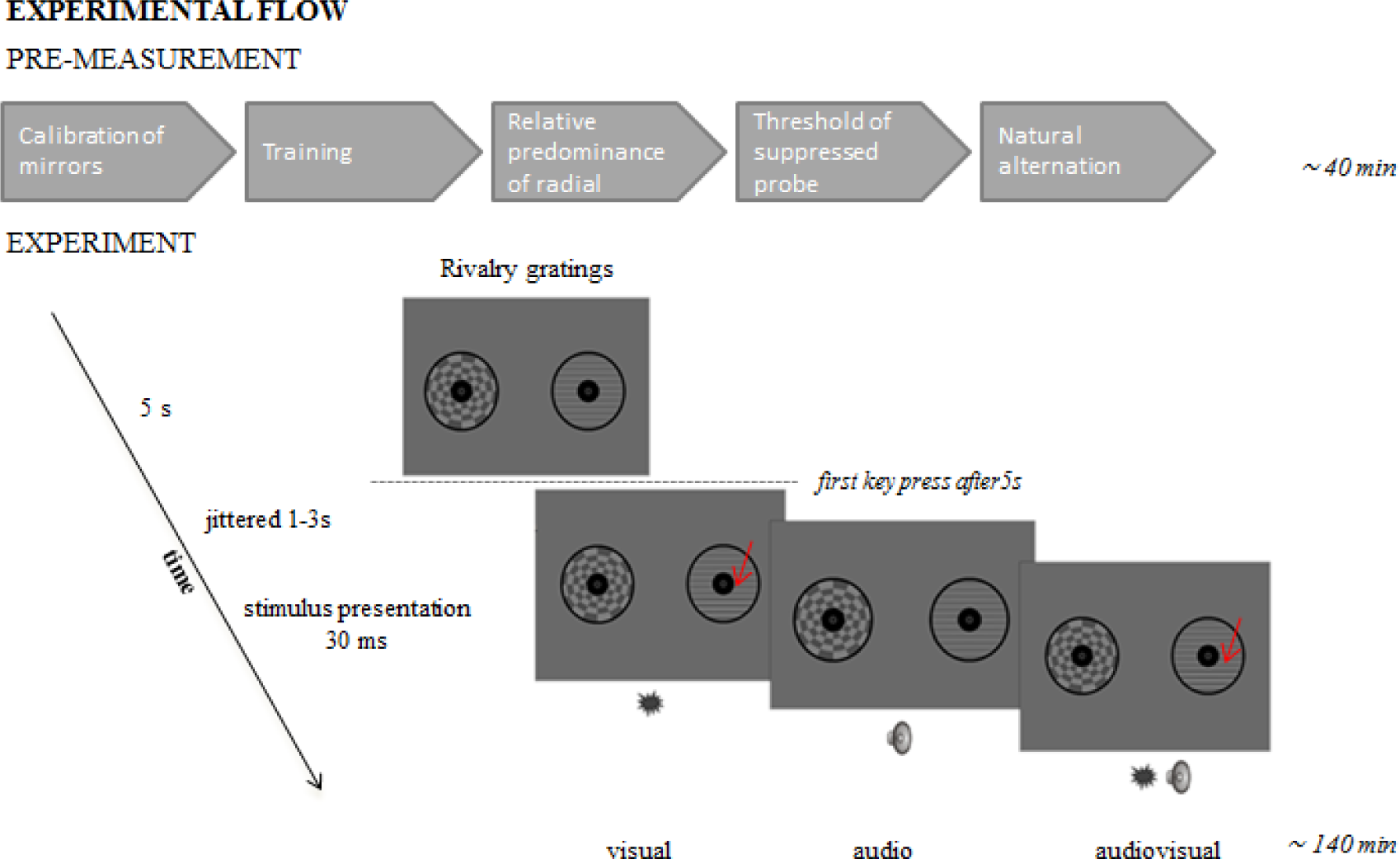
*Experimental flow.* Each session contained of two parts, starting with different steps of pre-measurement and followed by the experiment by itself. During the experiment each trial started with a 5-s-waiting-period. Furthermore, SOAs were fixed to jittered 1-3s from the first coming key-press occurred after the 5-s-waiting-period. Visual, audio or audiovisual events were presented on the dominant (D) or suppressed (S) Gabor patch. The flash was presented on the lower part of the gratings in visual and audiovisual conditions, marked by red arrows on the graph for better visualization.

## 4. Discussion

The focus of the current study was to seek for evidence supporting MSI based on bottom-up mechanisms alone. To do so, we used visual stimuli presented below awareness combined with sounds, in a protocol where top-down selection based on expectation or attention would play a minimal role. Thus, the question would be whether a sound can ‘rescue’ visual events to awareness by means of MSI resulting from bottom-up mechanisms. The behavioural results in the BR task revealed a cross-modal benefit, since the typical switch in perceptual state following a flash on the suppressed eye happened sooner when a sound coincided with the visual event. This would initially be consistent with the hypothesis of bottom-up MSI. However, this cross-modal benefit could not be univocally ascribed to bottom-up MSI (i.e., co-activation), because empirical data from audiovisual events did not deviate from the prediction of probability summation (PSM, e.g., postulated by Raab (1962)). This pattern of cross-modal facilitation without solid proof of MSI is very much in line with the previous behavioural study by Pápai and Salvador-Soto (Pápai & Soto-Faraco, 2017). Yet, because bottom-up MSI might have happened without surpassing the limit of probability summation in behaviour, we sought for neural correlates of audiovisual integration using EEG. According to the integration hypothesis, and the logic used in a multitude of previous studies (for above-threshold stimuli), if audiovisual ERPs surpass the threshold set by the summed unimodal ERPs, then MSI can be inferred (Fort et al., 2002a, 2002b; Giard & Peronnet, 1999; Molholm et al., 2002; Teder-Sälejärvi et al., 2002). Despite a sign of non-linear effect in a late time window (around ∼400ms post stimulus) when lowering the significance threshold, we failed to see evidence for bottom-up integration at sensory stages of processing. Since the ERP analyses could not confirm the bottom-up integration hypothesis, hence by default favoured the alternative hypothesis of the independent contribution from each unisensory response to the cross-modal ERP.

Before interpreting this result, it is relevant to discuss the neural correlates of the constituent, single modality, stimuli. Visual evoked potentials to events presented below awareness are often characterised by decreased amplitude of P1 (Kornmeier & Bach, 2005; Mathewson et al., 2009; Roeber et al., 2008), that can be even missing for threshold stimuli (Pins & Ffytche, 2003), as was the case in our study. In binocular rivalry paradigms, a diminished N1 and late positivity are often revealed for visual switching from suppression to awareness (Kaernbach et al., 1999; Roeber et al., 2008), though this pattern seems to be different when strong (very salient) visual probes are presented in the suppressed eye (Metzger et al., 2017; Valle-Inclán et al., 1999). Despite any of the previous BR paradigms measuring VEPs directly comparable to our set up, the VEPs for faint visual probes embedded in rivalry gratings seem to go in line with previous studies. In our data, both the N1 as well as a late positivity component were evoked in suppressed as well as dominant conditions. Although in both cases these visual components seemed to be attenuated under suppression, in accordance with previous BR studies measuring VEPs to perceptual switches (Kaernbach et al., 1999; Roeber et al., 2008), these effects were statistically unreliable, probably limited by the few number of trials/condition (since this analysis did not take part of the initial purpose of the study). Yet, what is important for the logic of interpretation in this context is that N1 as well as late positivity were effectively evoked by our visual stimuli even under suppression. Because these components have been also associated to attention capture and orienting respectively, their presence would confirm the possibility of attentional capture by the suppressed stimuli (Railo et al., 2011) (indeed, the small cross-modal ERP effect by the P300 time window would support that).

The auditory events in our study produced a prototypical Auditory Evoked Potential (AEP) with large response amplitude, given their above threshold strength, over parieto-occipital areas (Giard & Peronnet, 1999; Molholm et al., 2002). The audio stimulus was supra-threshold regardless of whether the target visual percept was suppressed or dominant, hence its associated AEP with the typical N1-P3 complex reflecting to sensory processing followed by attentional mechanisms did not vary between these two conditions.

Possibly as a consequence of the weak visual and more robust auditory responses audiovisual ERPs where mostly driven by the auditory response. What is interesting is that regardless the lack of a bottom-up multisensory ERP response, above and beyond the summed unimodal responses in early time window; these audiovisual events did produce a behavioural advantage over each of the unisensory events. This pattern reinforces the idea that such behavioural advantage was not based on bottom-up integration in sensory processing, and probably originated further down the stream of information processing. This interpretation might be strengthened by the small non-linear interaction appearing at late time window, when sensory processing is already quite predisposed to attentional influence. Furthermore, although we initially targeted occipital electrodes in order to pick up visual responses, the exploratory analysis across all scalp electrodes confirmed the ROI-centred findings.

Thus, all in all the results from the cross-modal ERP responses is in line with the behavioural results given by probability summation, favouring the independent contribution of the unisensory stimuli to the cross-modal behavioural benefit, and suggesting that there is no necessity to postulate an additional MSI mechanism to explain this cross-modal advantage. Of course, this does not mean that bottom-up MSI does not occur in other circumstances, but the present result has the implication that purely bottom-up mechanisms may not provide sufficient means for integrative operations below awareness. We tried to single out such mechanisms using audiovisual events where the visual component was presented below awareness, and in the absence of selective top-down attention or expectation about the moment of appearance or the particular feature content of the suppressed event or its association with the sounds. Of course, this does not preclude the possibility that MSI occurs under these circumstances, if it is guided by top-down mechanisms leading to expectation or selective attention.

In the ERP literature, multisensory effects have been already demonstrated by using abrupt audio and visual stimuli (Fort et al., 2002a, 2002b; Giard & Peronnet, 1999; Talsma & Woldorff, 2005; Teder-Sälejärvi et al., 2002) although since the events were presented above level of awareness, furthermore expectation/anticipation (Fort et al., 2002b; Giard & Peronnet, 1999; Talsma & Woldorff, 2005; Teder-Sälejärvi et al., 2002), or top-down attention (Talsma et al., 2007; Talsma & Woldorff, 2005) may have had a crucial influence on enabling MSI, evading the present research question. Indeed, when attention has been explicitly manipulated, it seems that multisensory benefits, even those arising from simple temporal coincidence, weaken (Talsma et al., 2007; Talsma & Woldorff, 2005). The present results go one step forward, and suggest that some form of top-down modulation might be needed to enable even the most rudimentary forms of stimulus-driven MSI integration.

Nevertheless, it must be mentioned that behavioural multisensory benefits have been demonstrated in the past for unconscious visual stimuli (Lunghi & Alais, 2013; Lunghi et al., 2014; Zhou et al., 2010), although many of these multisensory benefits were never tested against a probability summation baseline, leaving the possibility of more parsimonious explanations open. Additionally, beyond the lack of probability summation baseline, in many other studies selective attention and/or expectation was simply not controlled for (Aller, Giani, Conrad, Watanabe, & Noppeney, 2015; Alsius & Munhall, 2013), resulting in the possibility of a top-down facilitation of the multisensory effect below awareness. Despite these effects are interesting in themselves, they do not speak directly to bottom-up MSI integration.

Hence, amongst the wider context of literature addressing whether MSI can occur for unaware stimuli, the main conclusion to emerge from the present findings is that even though cross-modal benefits can appear behaviourally, and can indeed furnish the observer with an adaptive advantage over unisensory situations, these benefits may not be exclusively grounded on bottom-up mechanisms of sensory integration. Rather, we suggest that the behavioural benefit for cross-modal events is, more likely, based on the combination of bottom-up attentional capture of each unisensory stimuli individually. Despite bottom-up multisensory integration is indeed still a possibility, for now the individual contribution seems to be the most parsimonious explanation when the potential for top-down modulation is minimized.

## 5. Publication details

### 5.1. Data availability

The datasets generated during and/or analysed during the current study are available from the corresponding author on request.

### 5.2. Funding sources

This study was funded by a Ministerio de Educación, Cultura y Deporte, Ref. FPU/1206471 to MSP, and ERC Starting Grant, Ref. 263145, Spanish Ministry of Science and Innovation (PSI2016-75558-P), and Comissionat per a Universitats i Recerca del DIUE-Generalitat de Catalunya (2014SGR856) to SSF.

### 5.3. Author contribution statement

SSF and MSP designed the study, MSP conducted the experiment, SSF, MSP and MT analysed the data and SSF and MSP interpreted the data, and SSF, MT and MSP wrote the paper.

### 5.4. Additional information

Competing financial interest: The authors declare no competing financial interest.

